# SIMOFF: Discovering the metabolic objective of the cell

**DOI:** 10.64898/2026.06.18.733185

**Authors:** Rachel Gaffney, Magnus Rattray, Jean-Marc Schwartz, Kate Meeson

## Abstract

There are huge variations in metabolic complexity between the different kingdoms of life. Whilst it has been shown that some simple, unicellular organisms such as *E. coli* direct their energetic resources towards maximising proliferation, the metabolic goals of more complex organisms are unclear. This is an especially important topic for engineered organisms, such as Chinese Hamster Ovary (CHO) cells, that have been modified to produce therapeutically relevant compounds. This metabolic goal is reflected in the objective function of a constraint-based model (CBM) and has a direct impact on the metabolic flux distribution that is predicted using Flux Balance Analysis (FBA). However, there is no broadly applicable approach to infer this objective function from experimental data, to ensure CBMs represent real growth conditions. Here, we developed SIMOFF (SIMulated annealing Objective Function Finder) to infer the most appropriate objective function from minimal experimental flux data. Our applications of SIMOFF to *S. cerevisiae* demonstrated that the most suitable objective function is dependent on key metabolic phenotypes, even when the same organism and conditions are being modelled. Furthermore, we demonstrated the translatability of SIMOFF through application to CHO cells, where we showed that a SIMOFF-inferred objective function improved the accuracy of gene essentiality simulations, resulting in more reliable experimental target predictions.

## Introduction

Constraint-based models (CBMs) are computational reconstructions of cellular metabolism that allow us to predict experimental phenotypes. To personalise these models, sample-specific enzyme expression values are integrated with a species-specific reaction network. Constraint-based modelling has improved our understanding of biopharmaceutical production in mammalian cells (1). For example, CBMs are commonly used to estimate genetically engineered phenotypes through the simulation of gene deletions or overexpression. This approach has been combined with machine learning to study resource allocation following the overexpression of secretory pathway genes (2,3).

However, constraint-based models must be correctly parameterised, as inconsistencies in parameterisation can introduce significant uncertainty into model analyses (4). A key component of constraint-based modelling is the objective function, which defines one or more reactions to be maximised during linear programming to obtain a single solution in flux balance analysis (FBA) (5). The objective function is intended to represent the long-term evolutionary goal of the cell, and specifying one is essential because, without it, the system of equations would be underdetermined.

Some flux analysis methods, such as flux sampling, do not require an objective function (6). However, flux sampling is computationally expensive, especially when applied to thousands of gene engineering simulations. Furthermore, samples may not reflect a specific condition of interest without imposing further constraints. For the more computationally efficient FBA approach we must consider how to choose, or infer, the most appropriate objective function for the sample and condition of interest.

The appropriate choice of objective function depends on the organism and conditions being studied, and it should be directed by experimental measurements. In simpler organisms, such as *Escherichia coli*, common objective functions include the maximisation of biomass production (proliferation rate), ATP yield or maintenance energy, or the minimisation of redox potential (7). Similarly, in *Saccharomyces cerevisiae*, the maximisation of biomass production is a standard objective function, sometimes in combination with the minimisation of carbon dioxide, NADH or NADPH production (7).

The choice of objective function is more ambiguous for complex, mammalian cell types. A common cell type used in bioprocessing are Chinese Hamster Ovary (CHO) cells, which are the expression platform for the majority of humanised monoclonal antibodies (mAbs) (8). Precisely how we parameterise CHO CBMs and their objective function has important industrial implications.

During a fed-batch culture, the metabolic profile of CHO cells shifts as it adapts to the available culture medium and cell density. Initially, cells exhibit a higher growth rate, accompanied by glucose and glutamine consumption, and lactate and ammonium secretion (9). Then, the cells move into an exponential growth phase, where they switch to lactate consumption and the rate of mAb production almost doubles (10). Finally, as nutrients become limited, CHO cells enter the stationary phase, where the proliferation rate stagnates and mAb production is at its highest (10).

Evidently, CHO cells have a complex, shifting metabolism, that involves a trade-off between growth, mAb production and nutrient exchange. It has been suggested that the objective function is the greatest source of uncertainty in CHO modelling, and that the maximisation of biomass production leads to overestimated growth rates and an overactive metabolism (11). Instead, a multi-objective function should be used to simultaneously consider all industrially important phenotypes (12).

To improve the accuracy of CHO CBMs, studies have suggested several alternative objective functions, including: the inclusion of non-growth associated maintenance energy constraints (13), the minimisation of non-essential nutrient uptake rates (14) and the nonlinear maximisation of ATP generation, whilst minimising the sum of squared intracellular fluxes (15). In addition, alternative flux analysis methods exclude the objective function entirely, such as flux sampling and temporal expression-based analysis of metabolism (6,16).

To resolve this issue, several methods have been suggested that use data-driven approaches to infer the most appropriate objective function. One such method, inverse FBA (invFBA) traces back from ^13^C-labelled substrates to output a linear, weighted multi-objective function (17). Another method, the SCOOTI algorithm, employs a meta-regression learner model to assign weighted coefficients to 52 biomass-related metabolite reactions, using bulk and single-cell omics data (18). Although these existing methods provide an excellent basis for method development, they have some limitations for application to CHO. For example, without comprehensive ^13^C-labelled flux distributions, invFBA is limited to non-genome-scale stoichiometric models (17). In addition, because the SCOOTI algorithm fits an objective function directly to omics data, discrepancies between omics measurements and actual metabolic flux activity will be reflected in the output (18). Furthermore, the weightings of vector coefficients and the error calculation could be biased by the great range in magnitude of reaction fluxes.

In this study, we sought to develop a novel method for inferring the most appropriate objective function from experimental measurements in order to improve the accuracy of constraint-based modelling. We termed this method SIMulated annealing Objective Function Finder (SIMOFF). SIMOFF outputs the simplest possible single or linear, multi-objective function to satisfy a set of user-defined, qualitative, non-continuous flux behaviours (either –1, 0 or +1, depending on reversibility).

Initially, we demonstrate that the FBA flux distribution heavily depends on the objective function. Secondly, we develop SIMOFF for *Saccharomyces cerevisiae –* a simple organism with well-characterised metabolic priorities. Then, with application in CHO, we demonstrate the accuracy of a SIMOFF-inferred objective function over standard objective functions. Finally, we show that a SIMOFF-inferred objective function improves CHO gene essentiality predictions, with validation against a large, publicly available dataset (19).

Our work shows that the specific combination and weighting of reactions in a multi-objective function is important for the accuracy of CBMs. We have used SIMOFF to predict potential gene engineering targets in CHO, whilst demonstrating that SIMOFF is applicable to a broad range of organisms with diverse metabolic goals, without the need for extensive input data.

## Materials and Methods

### Input and validation datasets

Gene expression data and metabolite exchange data were sourced from industrial fed-batch bioreactors and publicly available datasets were sourced for *S. Cerevisiae* phenotypic measurements and CHO gene essentialities (**Table *1***).

**Table 1.** Details of input and validation datasets.

| Dataset | Source | Identifier |
| --- | --- | --- |
| Gene expression measurements (CHO; time-series; fed-batch) | Bulk RNA-seq (20) | ENA study: ERP189558 |
| Bioreactor measurements (CHO; time-series; fed-batch) | Lactate exchange, ammonia exchange, growth rate and IgG productivity measurements (20) | https://doi.org/10.1002/bit.28982 |
| Phenotypic measurements (*S. Cerevisiae*; slower-growing and faster growing conditions) | Growth rate, glucose, ethanol, glycerol and ammonium exchange (21) | https://doi.org/10.1038/s41467-022-28467-6 |
| Gene essentiality dataset (CHO) | CRISPR screen (19) | https://doi.org/10.1016/j.crmeth.2021.100062 |

### Software and algorithms

We have included a table for software and optimizers to improve reproducibility (STAR Methods-style **Table *2***) and on the associated GitHub repository, we have included a requirements file with all packages and versions used to generate data for this article.

**Table 2.**
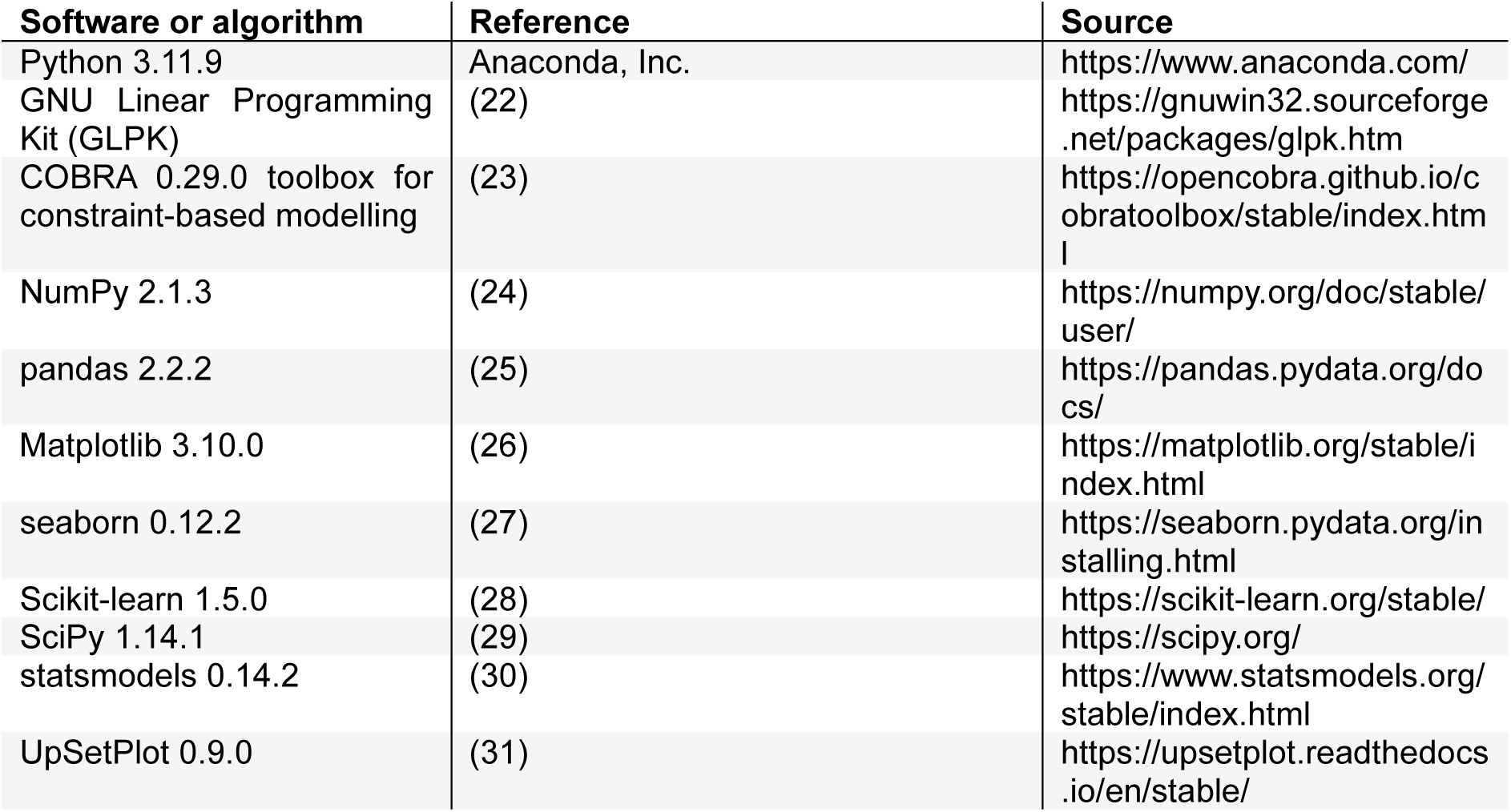
Details of software and algorithms.

### Construction of constraint-based models

To build the CHO constraint-based models, time-series transcriptomics was integrated into the iCHO2441 model (32), using methods detailed in a previous publication (20). Here, three models were constrained to represent the early exponential, late exponential and stationary/death CHO fed-batch culture phases. Gene expression measurements from samples taken at day 4 were integrated to constrain the early exponential model; samples taken at days 6, 7 and 8 to constrain the late exponential model; and days 11, 12 and 14 to constrain the stationary/death model. Originally, these models were validated using bioprocess measurements from key exchange reactions (20).

To build the yeast constraint-based models, minimal media conditions were calculated using the COBRA ‘minimal_medium’ function and applied to the consensus yeast genome-scale metabolic model: Yeast9 (23,33).

### Flux balance analysis

Flux balance analysis was performed using COBRApy (v0.29.0) (23). The GNU Linear Programming Kit (GLPK) (v5.0) was used to solve all linear optimisation problems (22).

To specify the objective function, the desired reaction vectors and coefficients were specified in a dictionary format and assigned to the ‘model.objective’ (**Table *3***).

**Table 3.** Objective function formulations. The objective functions that were specified across genome-scale models.

| Model | Objective function | Formulation |
| --- | --- | --- |
| iCHO2441<br>(Strain *et al.*,<br>2023) | Biomass maximisation | 'biomass_cho_prod': 1 |
|  | ATP (Complex V) maximisation | 'ATPS4m':1 |
|  | IgG maximisation | 'ICproduct_TRANSLATION_protein': 1 |
|  | Split objective (Figure 6b) | 'ICproduct_TRANSLATION_protein': 0.5,<br>'biomass_cho_prod': 0.5 |
|  | SIMOFF late exponential (Figure 6a) | 'ICproduct_TRANSLATION_protein': 0.9453,<br>'biomass_cho_prod': 0.0547 |
| Yeast-GEM<br>(Zhang *et al.*,<br>2024) | SIMOFF slower growing (Figure 4b and 4c) | 'r_2111': 0.771, 'r_1761': 0.005, 'r_1808': 0.015,<br>'r_1654': 0.210 |
|  | SIMOFF faster growing (Figure 4e) | 'r_2111': 0.95, 'r_1808': 0.05 |

### Gene deletion simulations

Gene deletion simulations were performed on the CHO late exponential constraint-based model. These were performed using the COBRA ‘single_gene_deletion’ function, which constrains the flux boundaries of all of the reactions that are associated with a specific gene to 0. A gene target was defined as one which upon deletion, reduced flux through the biomass reaction (‘biomass_cho_prod’). To further scrutinise potential targets, we filtered these essential genes to those that when deleted, also increased flux through the IgG productivity reaction (‘ICproduct_TRANSLATION_protein’).

To determine whether specific metabolic subsystems within iCHO2441 were enriched with target genes, a Fisher’s exact test was performed using the ‘multipletests’ function from statsmodels and the ‘hypergeom’ function from SciPy (29,30).

### SIMOFF algorithm

#### Implementation of SIMOFF

SIMOFF has been implemented as an open source Python package (https://github.com/katemeeson/SIMOFF) with example applications to yeast and CHO cells included, and with step-by-step instructions that can be adapted for any constraint-based model and input data.

The minimum necessary inputs for SIMOFF are a constraint-based model in SBML format, a list of input reactions, and a dictionary of qualitative criteria based on the experimental behaviour of an exchange reaction in the forward or reverse direction. For example, positive growth would be in the format ‘{Growth_ID: 1}’, where ‘Growth_ID’ is the model-specific ID for the biomass production reaction. The output of SIMOFF is a dictionary of the suggested objective function including reaction IDs and their coefficients, the maximum accuracy achieved (predicted versus measured fluxes), a dataframe of the predicted fluxes for each iteration, and a binarised agreement dataframe showing the mismatch of each iteration (0: disagree and 1: agree).

Python functions have also been included to generate the visualisations of the mismatch along the search (e.g. Figure 5c), how the coefficients change as the simulated annealing temperature decreases (e.g. Figure 6a) and the Upset plot to show SIMOFF maximum accuracy per input reaction combination (e.g. Figure 5b).

Corresponding code to generate each figure in this publication are included in the GitHub repo for the package.

#### Assumptions of SIMOFF

The SIMOFF algorithm is underpinned by FBA (5) and therefore assumes that the metabolic system is working at steady state (Sv=0), subject to flux capacity constraints *a*_i_ *<* v_i_ *< bi* and the objective function *Z*. To avoid infinite combinations of coefficients that could be output by SIMOFF, the sum of absolute value of coefficients in the linear multi-objective function must sum to one:

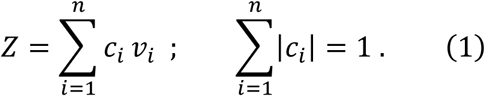

In SIMOFF, the user defines a list of several input reactions to be considered for a multi-objective function, assuming that these reactions contribute to the metabolic goal of the cell. These input reactions should represent the genetic engineering, immortalisation or known selective evolution of the cell line of interest.

Due to the formulation of the loss function, SIMOFF assumes that the absolute magnitude of a measured reaction flux is not relevant, only the directionality. SIMOFF compares the FBA-predicted and experimentally measured fluxes for the criteria reactions, which are converted to −1 (uptake), 0 (negligible exchange rates) or +1 (secretion) prior to input. This approach means that the magnitude of reaction fluxes does not skew the error measure, as it would if a continuous error measure, such as root mean square error, was used. This method aligns with omics integration algorithms that organise expression values into normalised ‘bins’ based on low, medium or high values and allows for error in the measurement of experimental values.

## Results

### The weighting of a multi-objective function controls FBA flux distribution

Initially, we wanted to determine how the composition of a multi-objective function controls the overall FBA flux distribution of a CHO CBM. The variables explored were the selection of reactions in the objective function and their weighted coefficients. We generated 500 randomly weighted multi-objective functions, containing six key bioprocess reactions (biomass production, IgG productivity, glutamine, glucose, lactate and/or ammonia exchange). Using these random multi-objective functions, CBMs were constrained for the early exponential, late exponential and stationary/death culture phases, using the iCHO2441 GEM (20,32). The resulting flux distributions were visualised using PCA and K-means clustering (**Error! Reference source not found.**) (elbow plots in ***Supplementary Figure 1***).

Firstly, we observed clustering across the flux distributions, which had been generated using the randomly sampled objectives (**Figure *1*a**, **Figure *1*b** and **Figure *1*c**). The optimal cluster frequency was unique to the culture phase, since there were three, six and five clusters that were formed across the early exponential, late exponential and stationary/death CHO models, respectively. Similarly, the individual reactions that were directing these clustering patterns were unique to the culture phase. For example, cluster separation along PC1 was primarily driven by the flux through potassium and creatine transport reactions (KHte, Kte, CREATte and CREATt) for the early exponential model (**Figure *1*a**); dopamine transport (DOPAt4_2_r and DOPAtu) and glycine transport (GLYt7_311_r and GLYSNAT5tc) for the late exponential model; and NAD/NADP redox reactions (DURAD2x and DURAD2) for the stationary/death model (**Figure *1*c**). However, regardless of the sample-specific transcriptomic constraints, the unanimous explanation was that the weighting of coefficients in a multi-objective does control the overall flux distribution.

**Figure 1.**
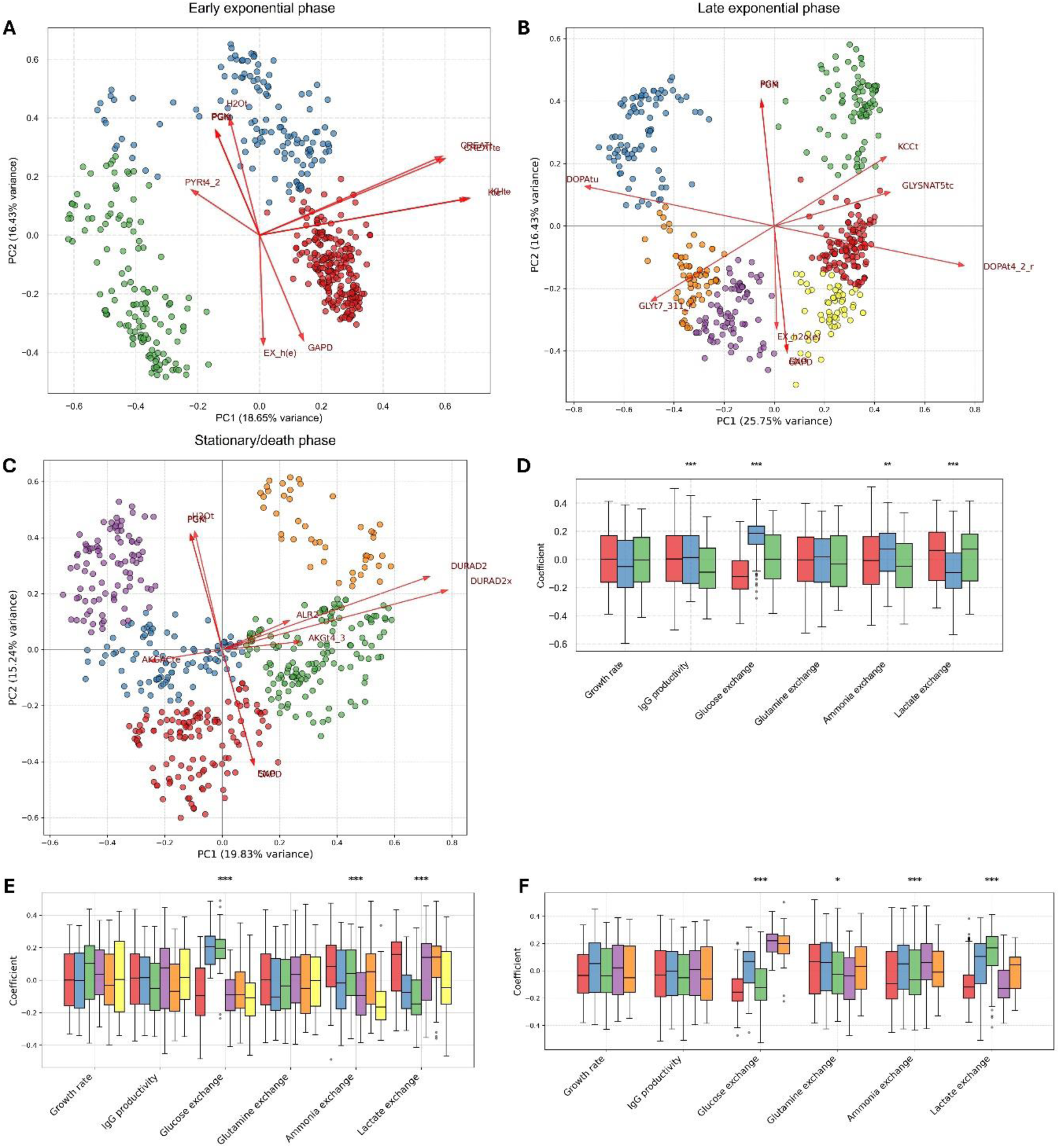
The weighting of a multi-objective function controls FBA flux distribution. **A.** PCA of flux balance analysis flux distributions using the early exponential phase CHO CBM. Solutions are *n*=500 random combinations of six reactions in a multi-objective function, including biomass production, IgG production, lactate, ammonia, glutamine and glucose exchange. Top five PC1 loadings and top 5 PC2 loadings have been annotated. **B.** PCA *n*=500 FBA flux distributions using the late exponential phase CHO CBM. Multi-objective parameters the same as in A. **C.** PCA *n*=500 FBA flux distributions using stationary/death phase constraint-based model. Multi-objective parameters the same as in A. **D.** Boxplot of the distribution of coefficients in the multi- objectives of *n*=500 FBA solutions, as described in A. Colour of box corresponds to cluster in A. **E.** Boxplot of the distribution of coefficients in the multi-objectives of FBA solutions, corresponding to C. **F.** Boxplot of the distribution of coefficients in the multi-objectives of FBA solutions, corresponding to E.

Another important question was whether some reactions in the multi-objective exert a greater influence on the clustering pattern of FBA flux distributions than others. In other words, did the optimisation of a particular reaction have a disproportionate effect on the overall flux distribution compared to other reactions? To study this, we plotted the distribution of coefficients in our six-part multi-objective function, across the distinct clusters that had been identified for each model (**Figure *1*d**, **Figure *1*e** and **Figure *1*f**). Here, we found that the weighted coefficients of glucose, ammonia and lactate exchange in the multi-objective function were significantly different across clusters for each of the three models. This shows that these three reactions have a particularly strong influence on the FBA flux distribution, relative to biomass synthesis (growth rate), IgG productivity and glutamine exchange.

This analysis showed that the objective function does impact the overall FBA flux distribution, and that specific reactions have a disproportionate effect on this flux distribution when they are optimised.

### The objective function influences model accuracy and gene engineering simulations

Once we had shown that the objective function affects the FBA flux distribution, we wanted to explore its effect on the accuracy of reaction flux predictions and the variability of gene engineering simulations. Here, we explored four key bioprocess reactions, that we had studied experimentally in previous work: biomass synthesis (growth rate), IgG productivity, lactate and ammonia production (20).

To study the accuracy of flux predictions, we plotted experimentally measured values metabolite exchange values against FBA predictions from the 500 randomly-sampled multi-objective functions described above (**Figure *2*a**, **Figure *2*b**, **Figure *2*c** and **Figure *2*d**). Of all four bioprocess reaction that were explored, the growth rate was the most inaccurately replicated by FBA and was underestimated by roughly 200-fold (**Figure *2*a**). On the other hand, some FBA solutions did accurately capture IgG productivity, ammonia and lactate production (**Figure *2*b**, **Figure *2*c** and **Figure *2*d**). In fact, the majority of flux predictions for lactate production in the late exponential and stationary/death models were around the experimentally measured values (**Figure *2*d**).

**Figure 2.**
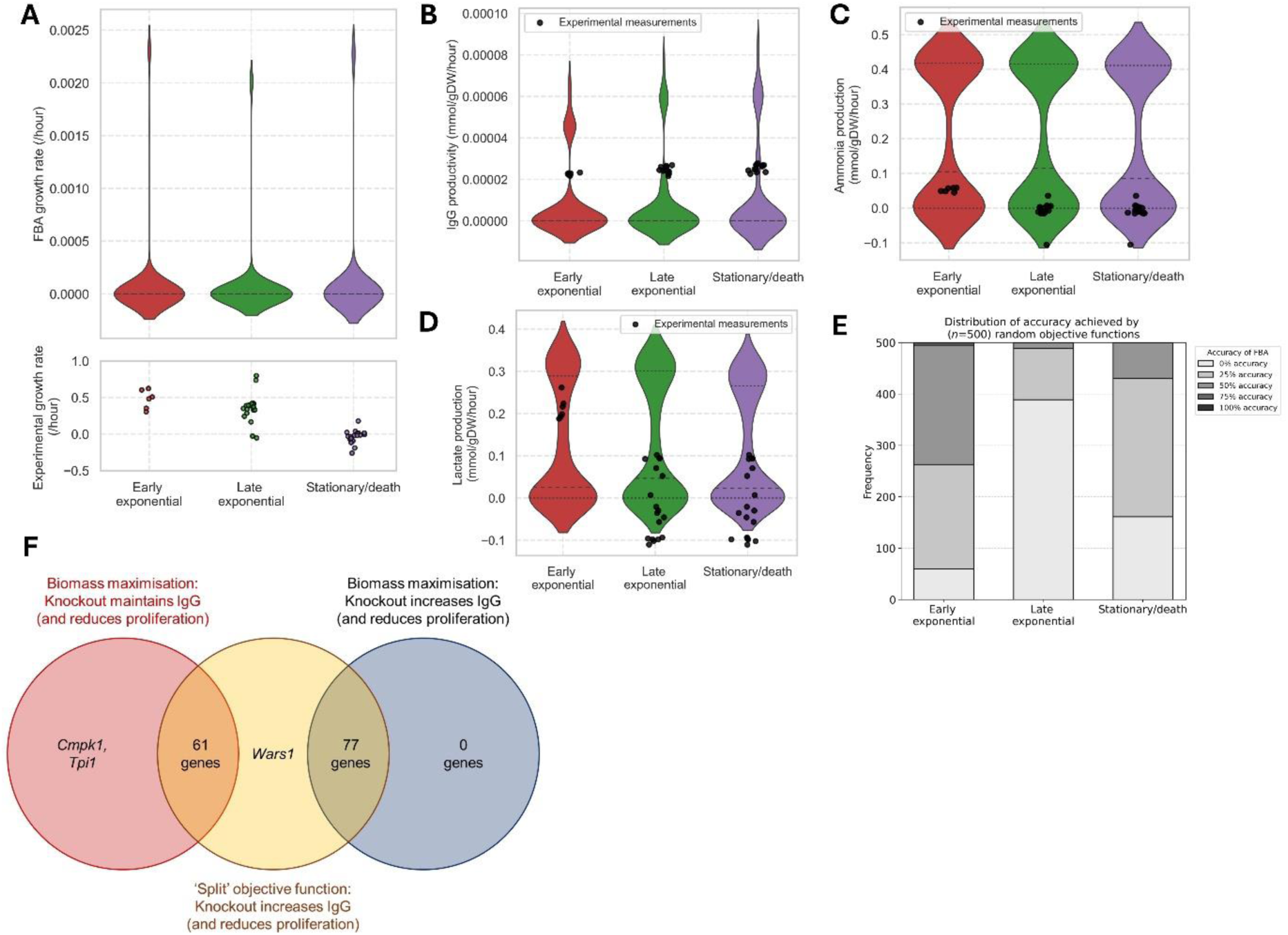
The objective function influences FBA accuracy and gene engineering simulations. **A.** Violin plot to show the distribution of growth rate predictions (hour^-1^) (*n*=500 with random multi-objective function) across the three culture phase CHO CBMs. Violin includes FBA predictions and individual plotted points are experimental values across several bioreactors. **B.** Violin plot to show the distribution of IgG productivity predictions (mmol/gDW/hour) (*n*=500) across three culture phase CHO CBMs. **C.** Violin plot to show the distribution of ammonia production predictions (mmol/gDW/hour) (*n*=500) across three culture phase CHO CBMs. **D.** Violin plot to show the distribution of lactate production predictions (mmol/gDW/hour) (*n*=500) across three culture phase CHO CBMs. **E.** Distribution of accuracy to a four-part, qualitative experimental criteria (bioreactor measurements of growth rate, IgG productivity, lactate and ammonia production either −1, 0 or 1 for consumption, no flux or production of metabolite). Accuracy has been calculated for *n*=500 FBA simulations, with random multi-objective functions across three culture phase-specific CHO CBMs. **F.** Venn diagram to show the intersection of potential gene target predictions, depending on objective function. Biomass maximisation, IgG maximisation and a 50:50 ‘split’ multi-objective function including both biomass and IgG maximisation were explored. Simulations ran on late exponential model, and a potential target was defined as a gene deletion that reduces growth rate (compared to mode) whilst maintaining or increasing IgG productivity. Wars1: Tryptophanyl-TRNA Synthetase 1; Tpi1: triose phosphate isomerase 1; Cmpk1: Cytidine/Uridine Monophosphate Kinase 1.

Although individually, FBA simulations could replicate most of these key bioprocess reaction fluxes, we wanted to know whether they were ever accurately predicted in combination with one another. From this analysis onwards, we chose to use qualitative reaction flux values for the experimentally measured and FBA fluxes, because without applying direct constraints, it is difficult to predict absolute values given uncertainties in biomass weight, mass balance of the biomass reaction and biomass composition (11). Therefore, we transformed absolute fluxes to discrete, qualitative values, describing the directionality of the metabolite exchange (−1, 0 or +1 for uptake, minimal exchange or metabolite secretion). Here, we found that across the three CHO models, no randomly sampled multi-objective function was able to parameterise a CBM that could correctly reproduce all four bioprocess reaction fluxes (**Figure *2*e**). The maximum accuracy reached by any objective function was 75%, which was achieved by only five of the 500 CBMs for the early exponential phase (**Figure *2*e**).

Gene engineering is an important application of CBMs, so we explored how varying the objective function formulation affected gene engineering simulations. For this analysis, we used the late exponential phase CHO model, since this is a popular point for growth arrest strategies that aim to redirect resources towards IgG production (34). We explored a full biomass maximisation, a 50:50 multi-objective maximising biomass and IgG with equal weighting and a full IgG productivity maximisation. A potential target was defined as one which maintains or increases IgG productivity, whilst reducing the rate of biomass synthesis, but maintaining a growth rate above zero.

Our gene engineering simulations showed that the objective function must include biomass synthesis maximisation to predict a non-zero growth rate, therefore, there were no targets predicted for the IgG maximisation objective function (**Figure *2*f**) (**Supplementary File 1**). In addition, when biomass synthesis was the sole objective, baseline IgG productivity was zero, so any gene deletion that increased it was considered a potential target. Furthermore, between the biomass maximisation and 50:50 split objective function, there was disagreement in how gene deletions affected IgG productivity. In particular, there were 61 genes (out of 139 overlapping targets) that were predicted to maintain IgG productivity using the biomass maximisation objective function but increase IgG productivity using the split objective function (**Figure *2*f**).

### Developing the SIMOFF package

These results show that whilst some CHO bioprocess behaviours can be accurately reproduced with randomly sampled objective functions, they cannot be all correctly predicted simultaneously. Furthermore, since the objective function directly influences flux predictions, this in turn controls gene engineering simulations. Since flux predictions and gene deletion simulations are key outputs of FBA, and often used for its validation, they should be embedded within the objective function selection process. These observations provided incentive for us to develop an algorithm for inferring the objective function from experimental measurements.

Our analyses show that the choice of objective function introduces significant uncertainty into FBA predictions. Individual reactions within the objective function can disproportionately impact the resulting flux distribution. Furthermore, the objective function strongly influences both the accuracy of flux predictions and the outcomes of gene engineering simulations. We demonstrated these effects across three CHO CBMs and, to address these challenges, we developed an algorithm to infer the most appropriate objective function from experimental data. This algorithm was termed ‘SIMOFF’ (SIMulated annealing Objective Function Finder), which we will now describe.

Our workflow uses simulated annealing to identify the globally optimum objective function of a CBM, through comparison of a qualitative experimental criteria to FBA predictions (**Figure *3*a**). Simulated annealing is a stochastic optimisation algorithm designed to identify global optima in complex, high-dimensional search spaces that may contain multiple local minima (35). Simulated annealing starts with a high ‘temperature’ that enables broad exploration of the solution space, gradually reducing this temperature to favour convergence toward an optimal solution (**Figure *3*a**).

**Figure 3.**
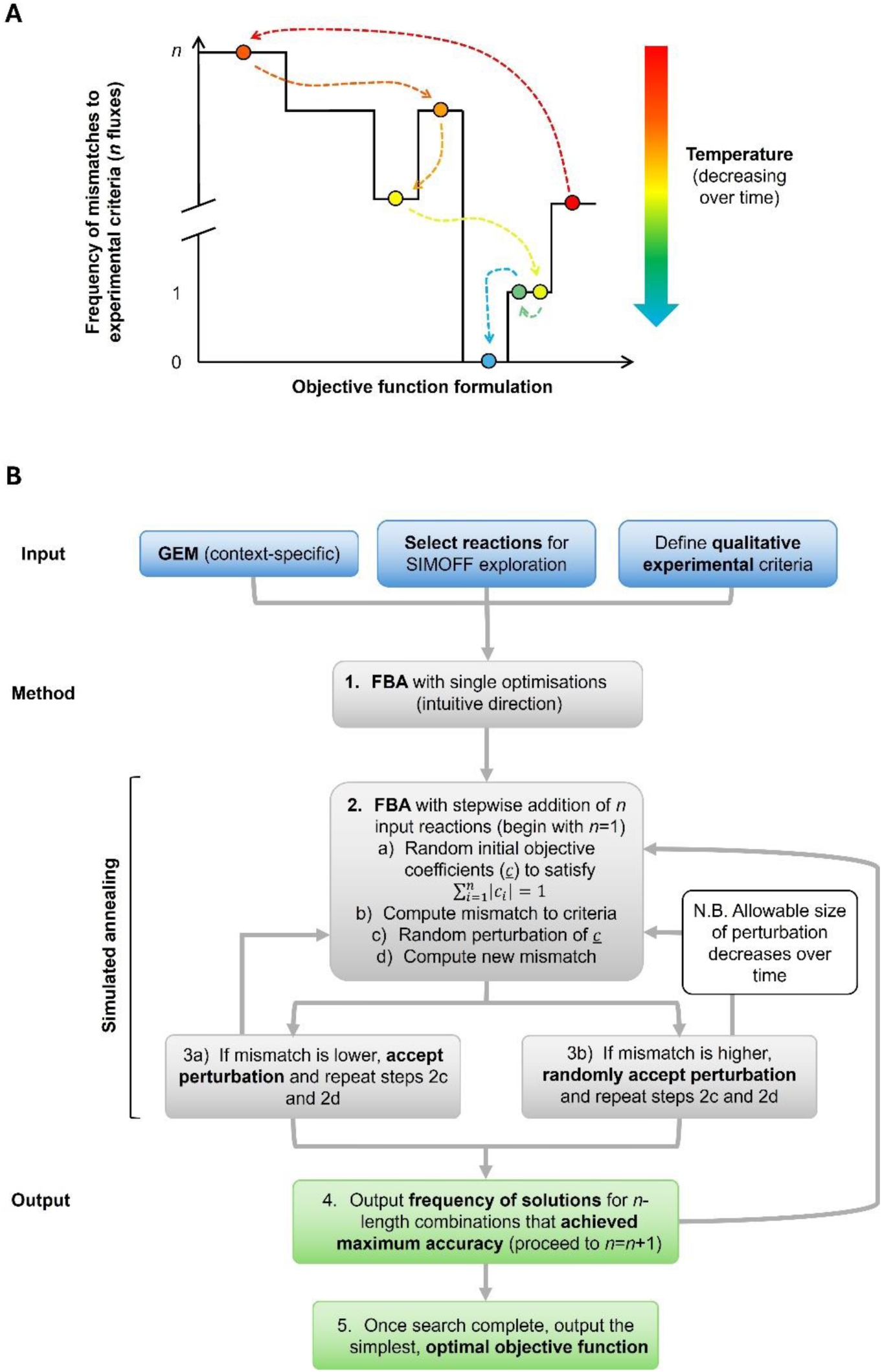
Workflow for SIMulated annealing Objective Function Finder (SIMOFF). **A.** Schematic for the process of simulated annealing to find the appropriate reaction coefficients for a multi-objective function, according to mismatch to a qualitative experimental criterion. Temperature decreases over time and fewer worse solutions are accepted. In a successful case, by the end of the search, a set of reaction coefficients that yield an FBA objective function with 100% accuracy (0 mismatches on y-axis) will be found. **B.** Workflow for SIMOFF approach. Input a context-specific GEM, a set of reactions to consider in the objective function and a qualitative experimental criterion (−1, 0 or +1 for fluxes, based on directionality). Proceed with SIMOFF, which incorporates simulated annealing and output the simplest objective function that best matches the experimental criteria.

**Figure 4.**
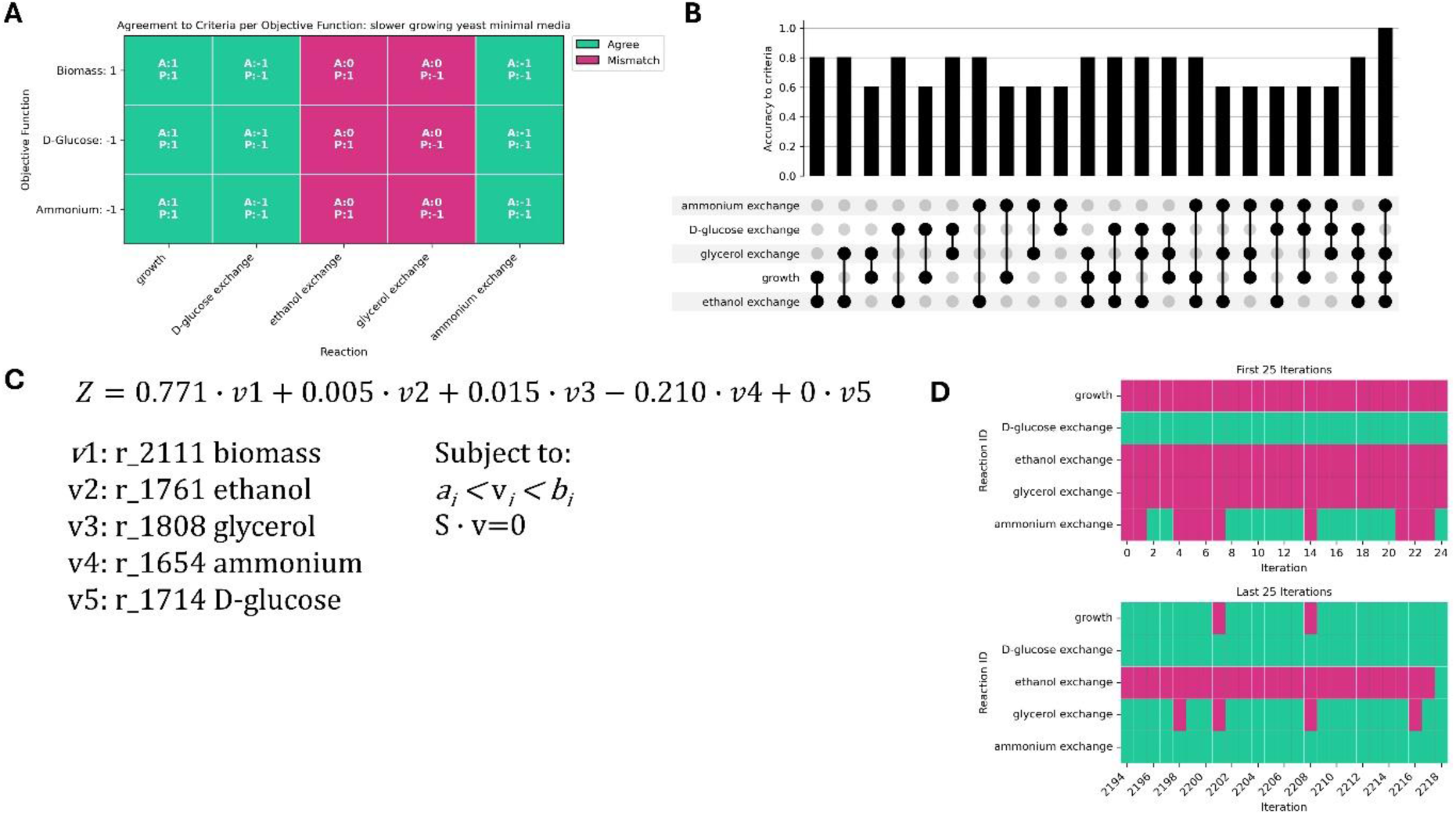
Application of SIMOFF to infer objective functions in slower growing *S. Cerevisiae* condition. **A.** Agreement between FBA predicted reaction fluxes (specifying the optimisation of single reactions) and experimentally measured fluxes (Elsemman *et al*., 2022) in the slower-growing yeast application. Teal indicates agreement between actual (A) and predicted (P) reaction fluxes, whilst pink indicates disagreement. Fluxes were transformed into qualitative values (−1: consumption, 0: zero flux or +1: secretion). The y-axis indicates the reaction in Yeast-GEM that has been optimised for (‘Biomass’: r_2111, ‘D-Glucose’: r_1714, ‘Ammonium’: r_1654’) and +1 or −1 coefficient refers to maximisation in either the forward or reverse direction, respectively. The x-axis indicates agreement between FBA and experimental values per each optimisation. **B.** Upset plot showing the results of SIMOFF searches in slower-growing yeast, including sequentially more reactions and exploring all reaction combinations in the objective (from input of 5 User-specified reactions). The y-axis indicates proportion agreement to experimental criterion and the search ends when an optimisation of 100% accuracy has been found. Dots on the x-axis indicates reactions that are being explored by SIMOFF for inclusion in the objective function. **C.** SIMOFF inferred multi-objective function for slower growing yeast condition. Subject to flux capacity constraints and the steady-state assumption. Using the following hyperparameters: max_iter=300, initialtemp=30 and maxfun=5000. **D.** Snapshot of the mismatch between FBA and experimental over the SIMOFF search for slower-growing yeast. Teal indicates agreement and pink disagreement. The x-axis indicates the iteration of the SIMOFF search, and the y-axis indicates agreement of specific FBA predicted reaction flux to corresponding experimentally measured value.

For the SIMOFF workflow (**Figure *3*b**), we chose to specify the input of a qualitative experimental flux criterion, where the values of −1, 0 or +1 (uptake, minimal exchange or metabolite secretion) are assigned to a metabolic flux, depending on reaction directionality and regardless of flux magnitude. Additional inputs include selected metabolic reactions to be explored within the objective function and a CBM. The loss function within SIMOFF is defined as the agreement (as a percentage) between the experimental criteria and FBA. So, for a four-part experimental criterion, this would look like 0%, 25%, 50%, 75% or 100% accuracy.

To begin with, SIMOFF will compare single optimisations of the input reactions to the experimental criteria (step 1; **Figure *3*b**), and then proceed to simulated annealing, including sequentially lengthier combinations of reactions and perturbing the reaction coefficients within the multi-objective function until the mismatch has been minimised and agreement is 100% (steps 2-4; **Figure *3*b**). Following the testing of all reaction combinations, the simplest objective function formulation that best fits the experimental criterion is output (step 5; **Figure *3*b**).

### Benchmarking SIMOFF on the well-characterised *S. cerevisiae* model organism

We chose to develop our method against *Saccharomyces cerevisiae* (baker’s yeast), since this is a well-characterised model organism with abundant publicly available data. Specifically, we chose to model a faster-growing (greater than 0.35 hour^-1^) and a slower-growing (less than 0.28 hour^-1^) condition, where the growth rate had been varied to mimic yeast response to sugar availability (Elsemman *et al*., 2022). These two growth rate conditions provided varied metabolic phenotypes, including: ethanol, glucose, ammonium and glycerol exchange. In this way, there were two separate qualitative experimental criteria from which our method could infer an appropriate objective function (**Error! Reference source not found.**).

For yeast simulations, the Yeast9 community-consensus genome-scale metabolic model was used (33) with minimal medium constraints. Five reactions were explored in the objective function (‘r_2111’: biomass, ‘r_1714’: glucose, ‘r_1761’: ethanol, ‘r_1808’: glycerol, ‘r_1654’: ammonium), and the flux distribution predicted by each optimisation was compared to an experimental criterion corresponding to the relevant growth condition (**Table *4***). From the five input reactions, there were 31 unique combinations of reactions to be explored within the objective function.

**Table 4.** Qualitative metabolic fluxes across yeast conditions. Qualitative experimental fluxes to form criterion for inferring the most appropriate objective function across slower-growing (less than 0.28 hour^-1^) and faster-growing (greater than 0.35^-1^) yeast conditions (*21*). Quantitative fluxes have been transformed into qualitative, using the following rules: −1: uptake, 0: minimal exchange measured and +1: secretion. Model IDs from the Yeast-GEM have been included for cross-referencing (*33*).

| Metabolite | Slower-growing condition | Faster-growing condition |
| --- | --- | --- |
| Biomass (r_2111) | 1 | 1 |
| Glucose (r_1714) | -1 | -1 |
| Ethanol (r_1761) | 0 | 1 |
| Glycerol (r_1808) | 0 | 1 |
| Ammonium (r_1654) | -1 | -1 |

Firstly, the slower growing yeast condition was explored. Prior to suggesting a multi-objective function, SIMOFF tested single optimisations of the input reactions. However, no single optimisation was able to achieve 100% accuracy to the experimental criterion. The single maximisation of biomass production, D-glucose uptake and ammonium achieved 60% accuracy, with each optimisation failing to accurately predict glycerol and ethanol exchange (**Error! Reference source not found.a**). The failure of single optimisations to predict experimental criteria indicated that a multi-objective function could be more appropriate.

The simplest multi-objective function that could achieve 100% accuracy specified the maximisation of biomass production, ethanol and glycerol secretion and ammonium uptake (**Error! Reference source not found.b**). Biomass production occupied the largest proportion (77%) of this multi-objective function, followed by 21% maximisation of glycerol uptake and negligible (<5%) proportions for ethanol and ammonium production (**Error! Reference source not found.c**). To understand the trajectory of the simulated annealing search, the mismatch to experimental criteria was plotted until the first instance of 100% accuracy was achieved (**Error! Reference source not found.d**). To begin with, when worse solutions could be accepted, growth, ethanol and glycerol exchange could not be resolved (**Error! Reference source not found.d**). As the search moved towards the global optimum, an appropriate multi-objective function was identified at 2,218 iterations, when ethanol exchange was finally resolved (**Error! Reference source not found.d**).

Once SIMOFF had resolved the slower growing yeast condition, we moved onto the faster growing condition. Here, the maximisation of biomass synthesis or D-glucose uptake was able to achieve 80% accuracy (**Figure *5*a**), indicating that this was a simpler problem than the slower growing yeast condition. Given that no single optimisation could achieve 100% accuracy, the SIMOFF search proceeded to consider multi-objective optimisation. SIMOFF identified a solution which reached 100% accuracy after 146 iterations (**Figure *5*b**) and this included the maximisation of biomass synthesis (95% of total objective function) and glycerol production (5%) (**Figure *5*c**). Earlier in the search, it was more difficult for SIMOFF to reproduce experimental biomass and ammonium exchange, but this was eventually resolved. In comparison with the slower-growing condition, biomass synthesis occupied a larger proportion of the total multi-objective function for the faster-growing condition.

We have benchmarked SIMOFF against a slower and faster growing yeast phenotype (21), to demonstrate its applicability across different experimental criteria. SIMOFF was able to identify multi-objective functions that could reach 100% accuracy to experimental criteria, although this was easier for the faster growing condition.

### Exploring the relationship between reactions in a multi-objective function in *S. cerevisiae*

We now wanted to study the relationship between reactions in a successful multi-objective function, including the variability of the coefficients. To allow this, we removed the exit statement in the SIMOFF algorithm to let the search proceed past the first instance of 100% accuracy.

First, we performed a new search for the faster growing yeast and SIMOFF output a multi-objective function including the maximisation of biomass synthesis and glycerol exchange (**Figure *6*a**). As the temperature decreased along the search, the coefficient of biomass production remained mostly positive, with some large jumps to very negative values, and the coefficient of glycerol exchange tended to zero (**Figure *6*a**). For this faster growing yeast condition, there was very little flexibility about the mean values of 0.945 (biomass synthesis) and 0.055 (glycerol exchange) (**Figure *6*b**).

For the slower growing condition, SIMOFF output a solution with 100% accuracy including biomass synthesis, ethanol, glycerol and ammonium exchange (**Figure *6*c**). Similarly to the faster growing yeast condition, the coefficient for the biomass synthesis reaction was the largest of all reaction vectors, with a range from 0.31 to 0.95 about the mean of 0.551 (**Figure *6*d**). The other key reaction for the slower growing condition objective function mean is the maximisation of ammonium uptake, with a mean coefficient of −0.435 and minimum and maximum allowable values of −0.68 and −0.025 **Figure *6*d**).

To visualise the SIMOFF trajectories, the flux distributions from the SIMOFF searches and (*n*=1000) randomly sampled objective functions were projected using PCA (**Figure *6*e** and **Figure *6*f**). For either growth condition, randomly sampled objective functions resulted in three distinct clusters. Top PCA loadings included reactions involving glycerol exchange and transport (r_1808, r_1172 and r_0489), which is unsurprising since glycerol exchange is specified in the SIMOFF output for both of these conditions. In addition, water diffusion reactions (r_1277 and r_2096) represented top PCA loadings for either condition. Ethanol and ammonium exchange were only specified in the SIMOFF output, and the presence of these reactions was reflected in the clustering pattern, where ethanol respiration, ammonium assimilation and subsequent ATP production (r_0438, r_0439, r_0226, r_1110 and r_1245) were amongst the top loadings (**Figure *6*f**).

In either case, the SIMOFF solutions of a lower accuracy (0-40%) sit broadly amongst all random sampling clusters (**Figure *6*e** and **Figure *6*f**). As the accuracy of the SIMOFF solutions increase to 100%, the flux distributions cluster more tightly, reflecting the smaller jumps being taken during the annealing search. In the PCA distribution for the faster growing condition, the 100% accurate flux distributions are distinct from the random samples, not occupying any cluster of flux distributions (**Figure *6*e**), which reflects the little variability in the 100% accurate flux distributions (**Figure *6*b**).

This analysis has shown that the flexibility and complexity of the most appropriate objective function depend on the experimental conditions that are being modelled. In addition, our work shows that there can be multiple possible ‘best’ objective functions, but the variability of these is also case-specific.

### A SIMOFF inferred objective function improves gene essentiality predictions in CHO

We have shown that SIMOFF improves the accuracy of *S. cerevisiae* CBMs, so next, we wanted to determine whether an experimentally inferred objective function could improve the accuracy of gene engineering simulations. This analysis also demonstrated the applicability of SIMOFF to a larger GEM with transcriptomic constraints.

Here, we utilised a CHO CBM that represents the exponential phase of a fed-batch culture. We modelled the exponential culture phase since there are distinct metabolic shifts that drive the transition from a proliferative to productive phenotype at this stage (10,36,37). Therefore, the most appropriate objective function is ambiguous. In addition, the exponential phase is a common point for growth arrest strategies that aim to redirect metabolic resources to increased productivity. The CHO CBM that we used has been previously published, and has matched transcriptomics and bioreactor measurements (growth rate, IgG productivity, lactate and ammonium exchange) (20).

To further benchmark SIMOFF, we evaluated the FBA performance of a SIMOFF inferred and standard objective functions. For this comparison, we included the standard biomass maximisation (38,39); IgG productivity maximisation (40); a 50:50 split objective function incorporating both biomass and IgG productivity, to explore whether the precise weighting of a multi-objective function is relevant; and ATP synthesis (Complex V) maximisation, because the inclusion of maintenance energy has been shown to improve CHO FBA (13).

SIMOFF output a multi-objective function for the exponential phase CHO CBM after only 13 runs (**Figure *7*a**). This objective function was dominated by IgG production maximisation (94.5%) with a small proportion of biomass maximisation (5.5%) (**Figure *7*a**). The SIMOFF-inferred multi-objective function was the only formulation that successfully reproduced the experimental criterion, outperforming all standard objective functions (**Figure *7*b**). This highlighted the importance of a multi-objective function for an exponential CHO CBM, since the biomass synthesis, IgG productivity and ATP generation single objective functions were not able to correctly represent ammonium uptake (**Figure *7*b**). In addition, this analysis signified the importance of the precise weighting of the coefficients in a multi-objective function, since the SIMOFF inferred formulation (94.5% IgG productivity maximisation and 5.5% biomass maximisation) was able to achieve 100% accuracy, yet the 50:50 split objective function maximising the same reactions was not (**Figure *7*b**).

**Figure 5.**
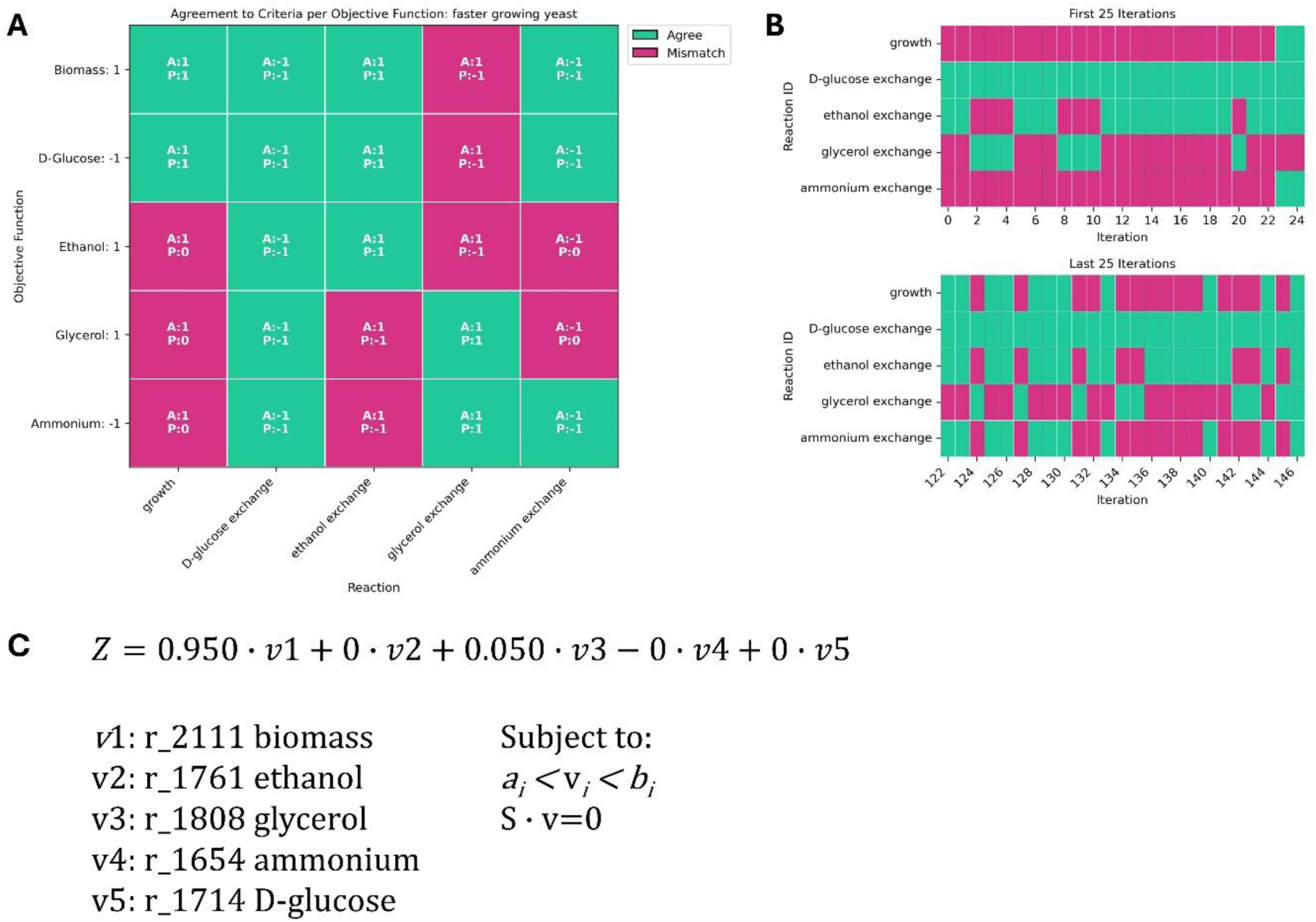
Application of SIMOFF to infer objective functions in faster growing *S. Cerevisiae* condition. **A.** Agreement between FBA predicted reaction fluxes and experimentally measured fluxes for the faster-growing yeast application. Teal indicates agreement between actual (A) and predicted (P) reaction fluxes, whilst pink indicates disagreement. Fluxes were transformed into qualitative values (−1: consumption, 0: zero flux or +1: secretion). The y-axis indicates the reaction in Yeast-GEM that has been optimised for. **B.** Snapshot of mismatch between FBA and experimental over the SIMOFF for the faster-growing yeast application. **C.** SIMOFF inferred multi-objective function for faster growing yeast condition. Subject to flux capacity constraints and the steady-state assumption. Using the following hyperparameters: max_iter=50, initialtemp=30 and maxfun=2000.

**Figure 6.**
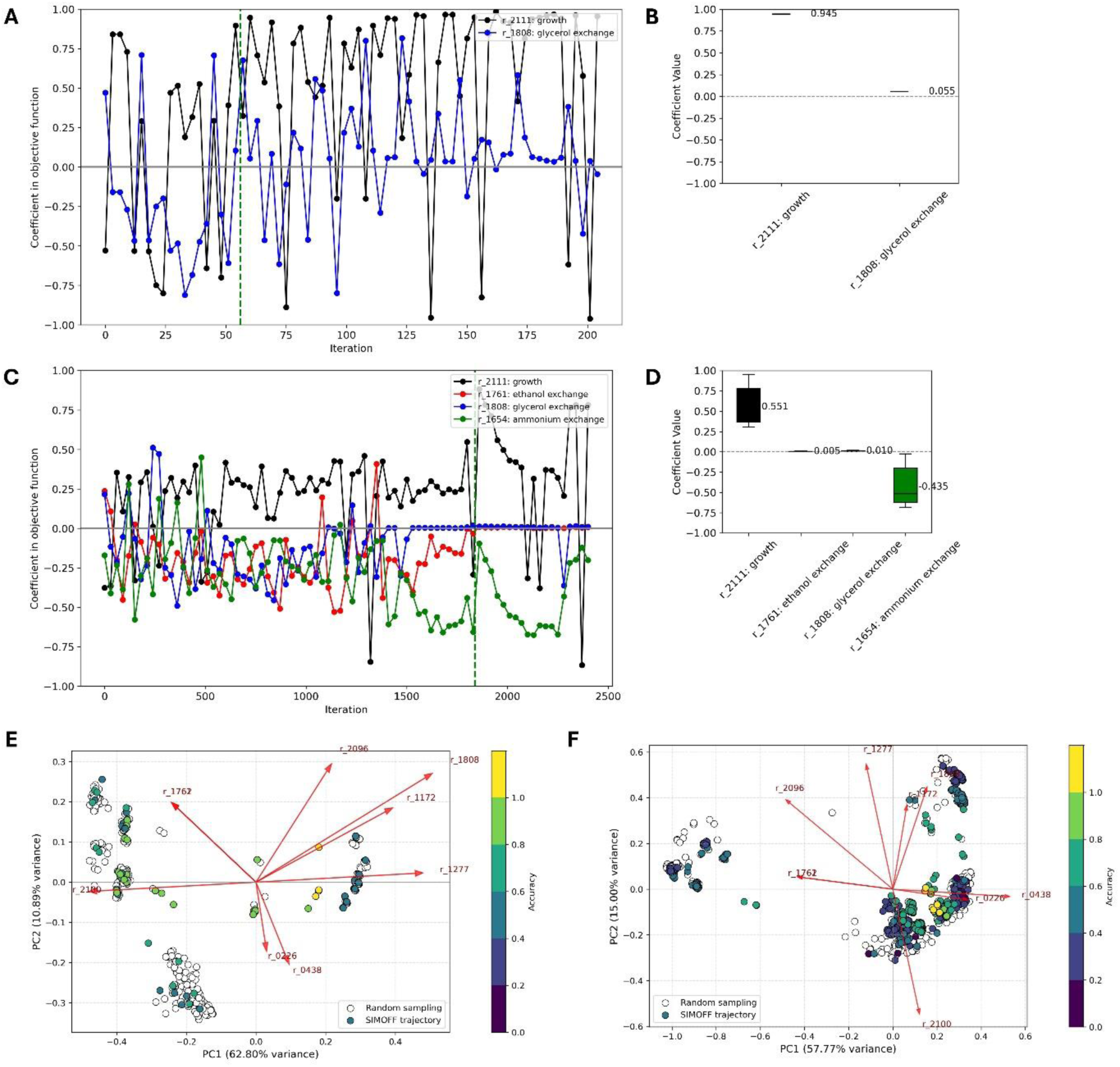
Exploring the relationship between reaction coefficients in the SIMOFF inferred multi-objective functions for S. Cerevisiae. **A.** The coefficients of biomass (‘r_2111’) and glycerol exchange (‘r_1808’) across the SIMOFF search, where the experimental criteria for faster growing yeast was specified. Parameters for this search were: initialtemp: 30; max_iter: 50; maxfun: 2000. Horizontal green line at x=56 was the first iteration that matched faster growing yeast experimental criteria with 100%. **B.** The range of coefficients of r_2111 and r_1808 in the (*n*=19) 100% accurate solutions from SIMOFF search for the faster growing yeast condition. **C.** The coefficients of biomass (‘r_2111’), ethanol (‘r_1761’), glycerol (‘r_1808’) and ammonium exchange (r_1654’) across the SIMOFF search, where the experimental criteria for slower growing yeast was specified. Parameters for this search were: initialtemp: 30; max_iter: 300; maxfun: 5000. Horizontal green line at x=1838 was the first iteration that matched slower growing yeast experimental criteria with 100%. **D.** The range of coefficients of r_2111, r_1761, r_1808 and r_1654 in the (*n*=415) 100% accurate solutions from SIMOFF search for the slower growing yeast condition. **E.** PCA of the entire flux distributions of (*n*=1000) random objective functions containing r_2111 and r_1808 and the (*n*=19) 100% accurate solutions from SIMOFF search. Colour of spot denotes accuracy to experimental criteria. The top 5 loadings for PC1 and PC2 have been annotated. **E.** PCA of the entire flux distributions of (*n*=1000) random objective functions containing r_2111 and r_1808 and the (*n*=19) 100% accurate solutions from the faster growing yeast SIMOFF search. Colour of spot denotes accuracy to experimental criteria. The top 5 loadings for PC1 and PC2 have been annotated. **F.** PCA of the entire flux distributions of (*n*=1000) random objective functions containing r_2111, r_1761, r_1808 and r_1654 and the (*n*=415) 100% accurate solutions from slower growing yeast SIMOFF search. Colour of spot denotes accuracy to experimental criteria. The top 5 loadings for PC1 and PC2 have been annotated.

**Figure 7.**
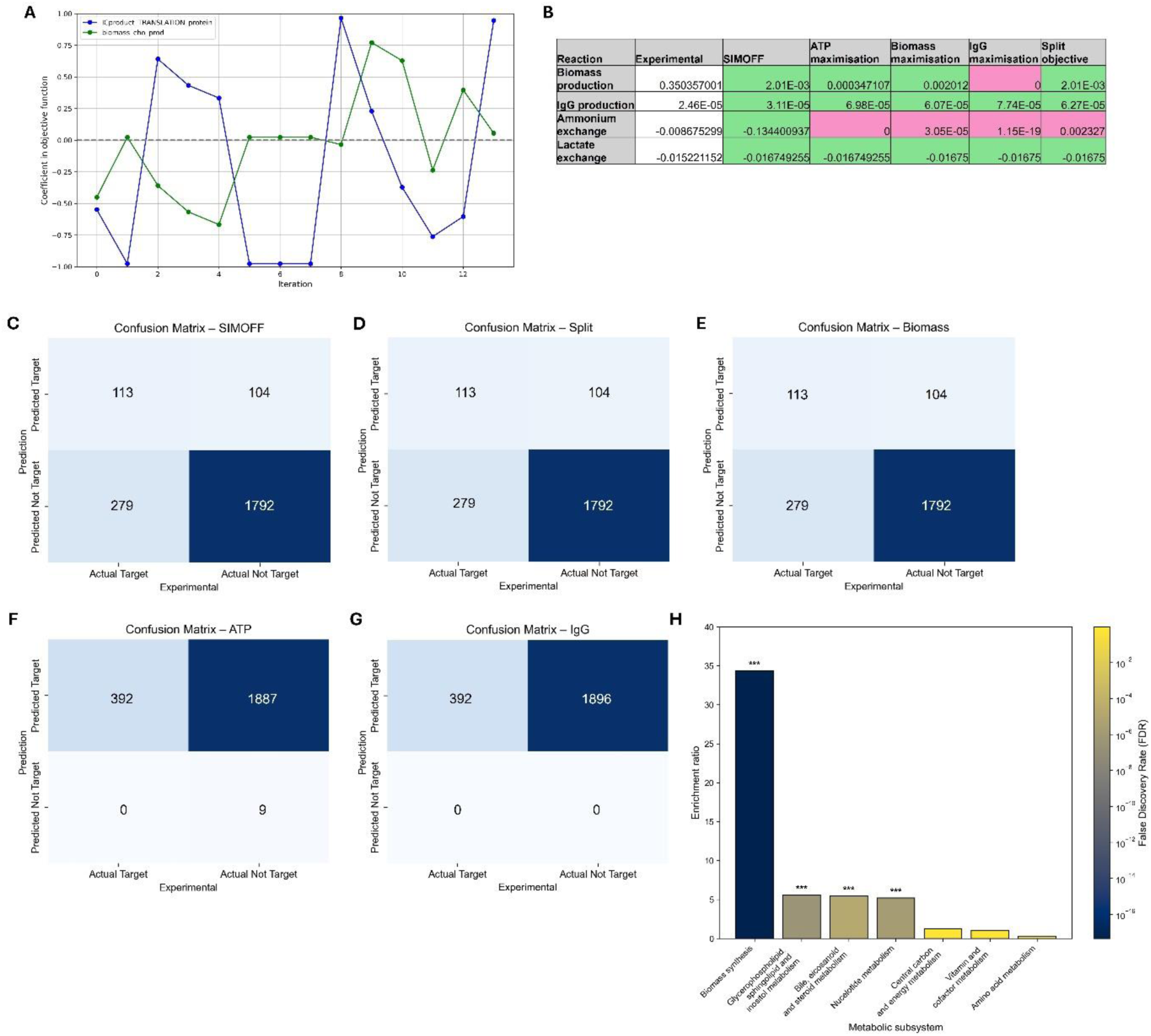
A SIMOFF inferred objective function improves the accuracy of direct flux predictions and gene essentiality simulations. **A.** The coefficients of the translation of IgG product (‘Icproduct_TRANSLATION_protein’) and biomass production (‘biomass_cho_prod’) across the SIMOFF search, where the experimental criteria for CHO DG44 cell line (growth, IgG productivity, lactate and ammonium exchange) at the late exponential phase in a fed-batch system has been specified. SIMOFF parameters for this search were: initialtemp: 300; max_iter: 50; maxfun: 5000. Search ended at iteration 13, where 100% accuracy to experimental criteria was found. The iCHO2441 model has been used with transcriptomics constraints. Optimal objective function was ’ICproduct_TRANSLATION_protein’: 0.9452826634320092 + ’biomass_cho_prod’: 0.0547173365679908. **B.** Flux balance analysis flux predictions using different objective functions: SIMOFF (from A), ATP production maximisation (Complex V; ‘ATPS4m’), biomass maximisation, IgG maximisation and a split objective function (1:1 ‘Icproduct_TRANSLATION_protein’: ‘biomass_cho_prod’ maximisation). Experimental predictions are three time points across six individual bioreactors (six biological replicates). **C.** Confusion matrix visualising the performance of the late exponential CHO model with the SIMOFF objective from A. Experimental data for CHO essential genes from previously published dataset by Xiong et al (PMID: 34935002). An essential gene in the model was defined as a decreased FBA growth rate after the gene deletion. **D.** Performance of a split objective function in the late exponential CHO model (1:1 ‘Icproduct_TRANSLATION_protein’: ‘biomass_cho_prod’ maximisation), relative to experimental dataset. **E.** Performance of biomass maximisation objective function in the late exponential CHO model, relative to experimental dataset. **F.** Performance of ATP production maximisation objective function in the late exponential CHO model, relative to experimental dataset. **G.** Performance of IgG maximisation objective function in the late exponential CHO model, relative to experimental dataset. **H.** Results of a Fisher’s exact test to show the enrichment of specific metabolic subsystems amongst gene targets. A target has been defined as a gene that when deleted in the late exponential CHO model, maintains a growth rate above zero, whilst improving IgG productivity relative to before the deletion. Using the annealing objective, there were 33 targets across the 7 subsystems visualised. False discovery rates (FDR) have been annotated (FDR<0.001 ***; FDR<0.01 **; FDR<0.05).

Next, to assess the performance of a SIMOFF inferred objective function in predicting essential genes, we validated gene deletion simulations against a publicly available dataset (19). This dataset was generated using a virus-free, genome-wide CRISPR screen and defined an essential gene as one which disrupted cell proliferation and viability following its knockout (19). There was 94% coverage of all model genes by this dataset.

Although absolute flux predictions depend on the objective function, there was an identical list of essential genes predicted by FBA when the SIMOFF inferred, biomass maximisation and split objective function were used (**Figure *7*c**, **Figure *7*d** and **Figure *7*e**). For these three objective functions, 83% of essentiality predictions (1,905 out of 2,288 total genes) were correctly predicted as either essential or not essential. Furthermore, these three objective functions achieved a specificity of 94.5% and a sensitivity of 28.8%, comparable to flux cone learning—a machine learning–based approach for predicting gene essentiality that does not require specification of an objective function (3). On the other hand, IgG maximisation and ATP generation maximisation showed poorer performance, wrongly characterising 82.5% of all genes as false negatives (a knockout was incorrectly predicted to not affect growth) and demonstrating 0 and 0.47% specificity (**Figure *7*f** and **Figure *7*g**).

Once we had validated our target lists against a publicly available dataset, we further defined a target as one which upon knockout would simultaneously reduce CHO cell growth and improve IgG productivity. Using the SIMOFF objective function, there were 33 potential experimental targets, using this definition. For the equally split, ATP generation maximisation, biomass maximisation objective functions, there were 21, 16, and 9 gene targets, respectively. Results suggest that maximising IgG productivity within the objective function likely overestimates the true flux, since no gene knockouts were predicted to increase the original, pre-knockout value. In fact, the closest FBA prediction for IgG productivity was obtained using the SIMOFF objective function (**Figure *7*b**).

Finally, we performed a subsystem enrichment analysis to identify which metabolic subsystems contained the highest proportion of gene targets (**Figure *7*h**). Here, we used the list of 33 gene targets that were predicted by SIMOFF (**Supplementary File 2**). Results showed that there was a statistically significant (*p*-value<0.05) overenrichment of targets in the ‘biomass synthesis’ (34-fold), with the majority of gene targets being genes encoding aminoacyl-tRNA synthetases. Potential gene targets were also significantly enriched across the glycerophospholipid, sphingolipid and inositol metabolism subsystem (5.6-fold enrichment) (*Cept1, Impa1, Agpat5, Cdipt, Cers2* and *Dgat1*); nucleotide metabolism (*Rrm1, Rrm2, Umps, Dhodh, Impdh1* and *Txnrd1*) (5.2-fold enrichment); bile, eicosanoid and steroid metabolism (*Nsdhl, Idi1, Fdps* and *Pmvk*) (5.4-fold enrichment).

## Discussion

Here, we have demonstrated that standard biomass objective functions are not always adequate to represent cellular metabolism with FBA. We tested a variety of alternative objective functions that have been suggested for CHO and found that these limit the ability of FBA to predict important metabolic phenotypes. In addition, these standard and alternative objectives lead to less accurate gene essentiality predictions. Therefore, we have developed SIMOFF to infer the objective function of a constraint-based model from experimental metabolite exchange data.

### A linear, multi-objective function represents realistic metabolic trade-offs

SIMOFF is designed to output a linear, multi-objective function, assuming that the cell makes a metabolic trade-off, where several conflicting metabolic goals are simultaneously optimised. This reduces the complexity of the solution, compared to, for example, a quadratic objective, and facilitates the biological interpretation of objective function in terms of a metabolic goal. If one reaction in the multi-objective function has a greater weighting than another, then one could interpret the optimisation of this reaction as being more central to the particular metabolic phenotype of the cell being modelled. CHO cells have been shown to optimise multiple, coupled biological processes, for example, productivity and product quality, which is crucial when considering experimental design (41). Recognised metabolic trade-offs have been reflected in several alternative objective functions that have been suggested for CHO constraint-based models, for example, maximisation of biomass production whilst minimising the uptake of essential nutrients (14). Non-linear objective functions have also been considered for CHO modelling, namely, the maximisation of fluxes through ATP-synthesising reactions with the minimisation of the sum of squared intracellular fluxes (15).

During testing, SIMOFF was applied against two slower and faster-growing yeast conditions (21). Experimentally, the slower and faster growing conditions have different growth rates (less than 0.28 h^-1^ and greater than 0.35 h^-1^), respectively) (21), but to satisfy the qualitative nature of the input criteria, these were both specified as +1 in SIMOFF. The objective function that SIMOFF suggested for the faster-growing yeastconsisted of 95% biomass production maximisation and 5% glycerol secretion maximisation. On the other hand, for the slower-growing phenotype, there was 77% biomass production maximisation,2.1% maximisation of ammonium uptake, 1.5% glycerol secretion and 0.5% ethanol secretion. These objective functions suggest that there is a greater prioritisation of biomass production in the faster-growing yeast condition.

The metabolic trade-off between growth and productivity that occurs at the exponential phase in CHO fed-batch culture was accurately captured by the SIMOFF objective function. SIMOFF suggested that there was a 94.5% maximisation of IgG productivity and a 5.5% maximisation of biomass production, showing that the emphasis was on IgG productivity. This is concordant with knowledge of this phase in culture, which is where the cell density if highest and the cells are at their most productive.

### Simulated annealing is a powerful and flexible method for fitting the objective function

Simulated annealing is an efficient and effective stochastic optimisation method, that we have shown to be capable of reaching the global optimum (100% accuracy) across both yeast and CHO case studies. Through the accepting of worse solutions during the gradual cooling of ‘temperature’, simulated annealing avoids becoming trapped in local minima (35). Furthermore, simulated annealing is an ideal method for this application owing to its simplicity (42).

SIMOFF has been designed to cater for minimal input data, without the requirement for extensive fluxomics and absolute flux measurements. The use of a qualitative criterion accounts for error and noise in measurements and means that there is no normalisation procedure required to balance the magnitude of flux measurements. This approach is similar to some of the discrete, relative methods for integrating omics data into constraint-based models; a systematic review showed that continuous or discrete values did not influence the accuracy of constraint-based models (43).

We defined the loss function as a mismatch to the experimental values (−1, 0 or +1 flux, compared to experimental). If we used a continuous error measure instead, such as Root Mean Squared Error (RMSE) as was employed by other tools (44), we would have to normalise the reaction fluxes.

In flux imbalance analysis, the concept of shadow prices demonstrated that some reactions showed greater variability in flux than others, but that this variation was not directly proportional to a change in objective value (45). In other words, a small change in flux can be biologically more significant for one reaction than for another. This means that if we were to allow a large difference between experimental and predicted value in SIMOFF to contribute disproportionately to the error measure, then the error measure might not be representative of actual biological signals. Therefore, in the SIMOFF loss function, we are focusing only on directionality of flux rather than its magnitude.

### SIMOFF identifies experimentally validated gene knockdown targets

In our late exponential CHO model, the SIMOFF-inferred multi-objective function gave more accurate gene essentiality predictions than other objective functions that have been proposed for CHO, namely ATP maximisation through Complex V and IgG productivity maximisation. Furthermore, the SIMOFF-inferred objective was the only objective function, out of several comparisons, that was able to predict all features of our experimental criteria.

Using our SIMOFF-inferred objective function, we predicted that there were 33 genes that when knocked out, would improve CHO IgG productivity by between 20-25%. The majority of these genes (12 out of 33) catalysed reactions within the biomass synthesis metabolic subsystem. The trade-off between proliferation and productivity in CHO cells is well acknowledged (46). Over the course of a fed-batch culture, cells move from an exponential growth phase to a stationary phase of high IgG productivity, which is underpinned by metabolic rewiring (47,48). Therefore, a common approach to improve IgG titre is to induce growth arrest at the exponential phase to redirect resources towards IgG production, through cell cycle arrest, gene engineering or temperature control strategies (49–52).

We identified 11 potential gene targets related to biomass production that when knocked out would improve CHO productivity. These targets included various aminoacyl-tRNA synthetases and research has shown that the selective expression of certain aminoacyl-tRNA synthetases enhances the modification of mAbs (53). The specific genes that our model has suggested should be knocked down could therefore shift the transcriptional machinery towards the more favourable aminoacyl-tRNA synthetase genes, as well as shifting metabolism towards productivity rather than proliferation.

The genes that were predicted to be essential for CHO proliferation, using our SIMOFF inferred objective function, were validated against the Xiong *et al*. dataset (19). We were able to independently validate our 33 top targets using a dataset by Marzluf *et al*. (54), where 20 of these targets were either listed directly, or their protein isoforms were described, as essential for CHO cell proliferation.

Lipid metabolism was a subsystem that was over-enriched with our predicted CHO targets, and this has been recognised as a worthwhile engineering target, where cells exhibit improved productivity via endoplasmic reticulum expansion (55). Our model predicted that silencing of the *Cers2* gene, which catalyses the synthesis of very-long-chain ceramides, could improve protein productivity. Validating this finding, there is evidence in CHO cells that mitosRNA or siRNA-induced knockdown improves specific productivity by approximately 60% (56).

We identified nucleotide metabolism as a metabolic subsystem that was enriched with potential gene targets. In particular, we suggested that *Rrm1* and *Rrm2* be considered as knockdown targets, and these genes are known to be associated with CHO cell proliferation (57). Therefore, the aforementioned metabolic trade-off between proliferation and productivity explains these targets.

Although we simulated the phenotypes of single gene deletions, these same genes could be candidates for a multi-gene engineering system. Our model predicted that the knockdown of *Dhodh*, *Umps* and *Impdh1*, all of which are involved in nucleotide metabolism, could improve productivity. Notably, these three genes contribute to a 10-gene rescue cassette that has been shown to improve CHO cell productivity (58). The fact that we could predict three of these ten genes supports that the objective function inferred by SIMOFF pleads to realistic simulations for gene engineering. In addition, it evidences nucleotide metabolism as a subsystem that should be further explored for gene targets.

### Limitations

Although SIMOFF was able to infer objective functions for the case studies mentioned here, as with all novel algorithms it will require further testing and benchmarking. For example, it would be interesting to apply SIMOFF to an *E. coli* study, since there is evidence that a multi-objective function is appropriate for *E. coli* models (59–61), especially if the modelling work involves engineered strains. We were able to test a broad set of conditions, including unicellular and multicellular organisms; omics constrained and non-omics constrained models; differently sized genome-scale models; four and five experimental criteria; and different growth rate conditions within the same model.

On the other hand, it is possible that simple, non-engineered organisms could be well-represented using a single biomass maximisation and that a multi-objective function is an overcomplication. We have accounted for this in our algorithm design by implementing a break statement that tests single optimisations before progressing to multi-objective optimisation.

SIMOFF uses simulated annealing to reliably identify the global optimum. However, as an increasing number of input reactions are specified by the user, the possible combinations of reactions to be explored within a multi-objective function will increase exponentially. In turn, the computational power required to run SIMOFF will also increase. Although we ran our simulations locally, and did not explore more than five input reactions, this limitation could be overcome by a high-performance computing cluster. Furthermore, alternative optimisation algorithms, such as Bayesian optimisation, which builds surrogate models from experimental data, could be used in place of simulated annealing.

Within SIMOFF, we incorporated the dual annealing optimisation from SciPy (29), which combines classical and fast simulated annealing. Although simulated annealing is theoretically guaranteed to find the global optimum, the algorithm has a slow convergence rate, which depends on a very gradual cooling schedule (62). However, variations of simulated annealing exist to overcome this, including the dual annealing method that we have integrated within SIMOFF and others, such as curious simulated annealing (63), that incorporate fast simulated annealing for improved convergence.

Beyond methodological limitations, an overarching challenge for all modelling studies is the incomplete nature of our current biological knowledge. In particular, the GEMs used in this work, and the pathways they contain, are restricted to reactions for which species-specific evidence exists or that have been inferred through homology with similar organisms. As a result, as additional metabolic pathways are discovered, there will be alternative routes to optimise a particular objective function, and the SIMOFF solution will change. Our simulations employed the most up-to-date consensus GEMs (32,33) and as new GEMs are reconstructed and released, users can easily substitute these GEMs within SIMOFF.

### Concluding remark

SIMOFF leverages sparse experimental data to infer a linear, multi-objective function, which reduces the uncertainty of FBA, by improving direct flux predictions and improving the accuracy of gene engineering simulations. Incorporating simulated annealing, we have developed SIMOFF, which can be applied to any biological system for which there is experimental metabolic flux data. This work is not restricted to flux predictions and gene engineering simulations but could infer the metabolic goal of unknown cell types, given limited experimental measurements. We have shown that a multi-objective function is an appropriate way to represent the metabolic goal of a secretory CHO cell. Using this objective, we identified several gene targets for knockdown that are predicted to redirect cellular resources towards increased IgG productivity. Encouragingly, some of these targets have already been explored experimentally, supporting further investigation into the remaining targets and positioning SIMOFF as a valuable tool to improve the accuracy of constraint-based bioprocess modelling.

## Acknowledgements

The authors acknowledge financial support from a Prosperity Partnership grant (EP/V038095/1) funded by EPSRC, BBSRC and FUJIFILM Diosynth Biotechnologies. This grant involved the University of Manchester, University of Edinburgh, University of York and staff at FUJIFILM Diosynth Biotechnologies. We acknowledge the Edinburgh Genomics department for transcriptomics data and analyses. We thank the many researchers from all collaborating universities for scientific discussions.

## Supplementary material

- **Supplementary File 1:** gene knockout simulations on CHO late exponential model, with biomass, IgG and split objective function. Contains categorical (‘increased’, ‘same’ or ‘decreased’) classifications for how a gene knockout affects biomass synthesis and IgG productivity and absolute fluxes from FBA.
- **Supplementary File 2:** gene knockout simulations on CHO late exponential model, using a SIMOFF inferred objective function. Targets have been split into metabolic subsystems after a subsystem enrichment analysis.

