## Supplementary figures for "SIMOFF: Discovering the metabolic objective of the cell"

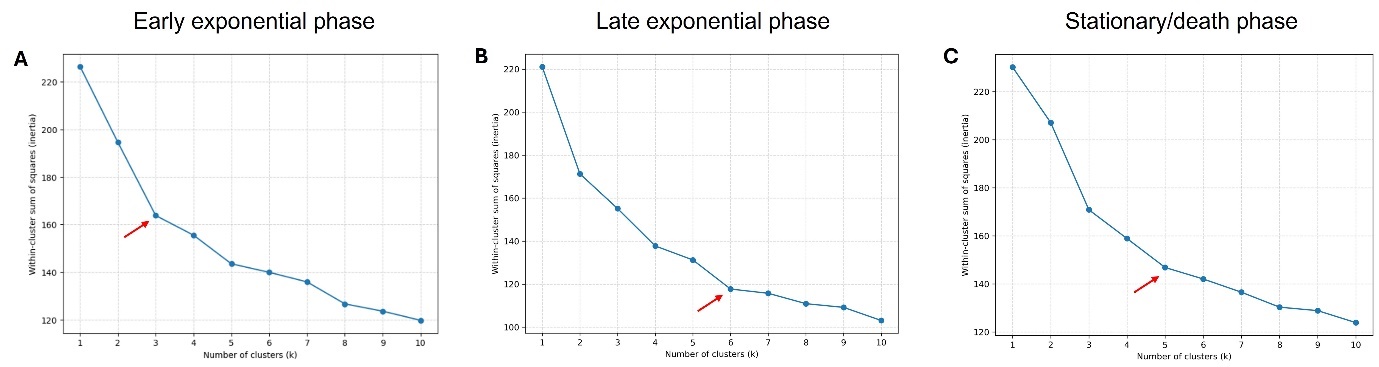

**Figure 1. Elbow plots for optimal cluster frequency calculations, for Figure 1.**

**A.** Elbow plot to calculate the optimal cluster frequency for (*n*=500) FBA simulations using an early exponential phase constraint-based model. Solutions are *n*=500 random combinations of six reactions in a multi-objective function, including biomass production, IgG production, lactate, ammonia, glutamine and glucose exchange. The absolute sum of the coefficients of this multi-objective must be 1 (negatives, decimal values and zeros are allowed). Optimal cluster frequency has been indicated on plot. **B.** Elbow plot for late exponential phase constraint-based model, same simulation parameters as in A. **C.** Elbow plot for stationary phase constraint-based model, same simulation parameters as in A.
