## Supplementary figures and images for "SIMOFF: Discovering the metabolic objective of the cell"

### Supplementary figure 1

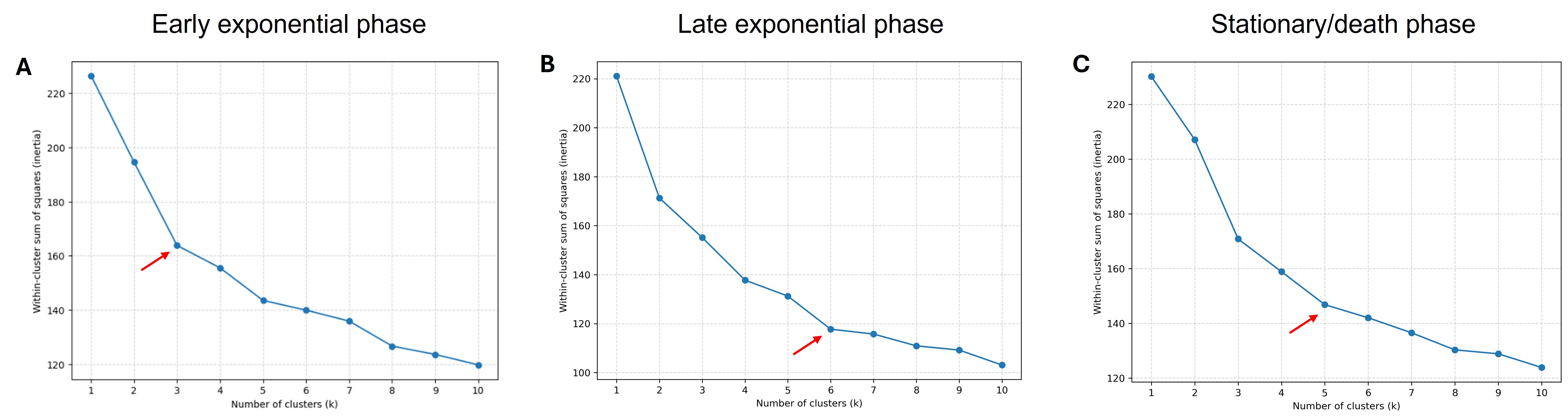
